# Differentiating Causes of Post-Surgical Conductive Hearing Loss with Optical Coherence Tomography Vibrometry

**DOI:** 10.64898/2026.01.10.698837

**Authors:** Xiaojie Yang, David P. Morris, Robert B. A. Adamson

## Abstract

**Purpose:** To evaluate whether optical coherence tomography (OCT) vibrometry can differentiate causes of conductive hearing loss (CHL) following ossiculoplasty, specifically distinguishing among effusion, soft-tissue fixation, and prosthesis disconnection.

**Methods:** We simulated three post-surgical CHL conditions, effusion, soft-tissue fixation, and prosthesis disconnection, in a cadaveric temporal bone model (*N* = 10 per condition), with a partial ossicular replacement prosthesis (PORP) and cartilage graft implanted via a retrotympanic approach. A custom-built swept-source middle-ear OCT system (*λ*_0_ = 1550 nm, Δ*λ* = 40 nm) was used to capture cross-sectional images and mobility measurements at the umbo and center of the cartilage graft across six stimulus frequencies (500, 750, 1000, 1500, 2000, and 3000 Hz). Mobility values (point velocity normalized to sound pressure) served as input features for a Random Forest classifier. Changes in mobility from baseline were also statistically analyzed.

**Results:** The classifier achieved 90.9% accuracy (40/44 datasets) in differentiating conditions in leave-one-out cross-validation, and 100% when trained on the full dataset. Simulated effusion and soft-tissue fixation were associated with broadband mobility decreases of 17.5 dB and 8.0 dB, respectively, averaged across both locations. Prosthesis disconnection at the stapes and at the tympanic membrane led to low-frequency (500 and 750 Hz) mobility increases of 9.9 dB and 11.1 dB, respectively, averaged across both locations.

**Conclusion:** In a cadaveric model of post-surgical CHL, OCT vibrometry accurately distinguished effusion, soft-tissue fixation, and prosthesis disconnection. The ability to identify the cause of post-surgical CHL highlights OCT vibrometry’s potential to assist clinical decision-making in revision surgery.

## Introduction

Optical coherence tomography (OCT) is emerging as a promising new tool for middle-ear diagnostics [1]. OCT has been explored by multiple groups for its use in imaging the middle ear [1–7]. The ability of OCT to provide simultaneous two-dimensional (2D) and three-dimensional (3D) images of middle-ear anatomy [4–7] and co-registered, spatially resolved vibration measurements of middle-ear structures in response to sound [6–8] makes this technology an attractive tool for the investigation of conductive hearing loss (CHL). OCT offers potentially powerful diagnostic information regarding the acoustic mobility of middle-ear structures that is currently unavailable with standard-of-care technologies [1–2].

Early studies into OCT of the ear have explored a wide range of potential applications, including measuring tympanic-membrane (TM) thickness [9], identifying dimeric TM segments [3] and cholesteatomas [10], assessing the post-operative status of middle-ear implants [6,11–12], differentiating various types of otitis media [2,13–14], and detecting cochlear implant migration [15]. In the present study, we investigate the application of OCT imaging and vibrometry to a new clinical question: the differentiation of various causes of CHL that recurs following middle-ear surgery. Studies of long-term hearing outcomes following ossiculoplasty indicate that initial hearing results are poor or that CHL returns post-surgically in about 10-15% of cases [16–17]. With current diagnostic methods, it can be difficult to distinguish various reasons for such failures and, hence, to decide how to manage these cases. Implants sometimes fall out of position, a problem that can generally be corrected with revision surgery. Middle-ear effusion can also cause a return of hearing loss following surgery. However, it is difficult to diagnose middle-ear effusion in a post-surgical ear because a significant portion of the TM is usually covered by an optically opaque cartilage graft, making it difficult to visualize fluid or to interpret tympanometry measurements. In contrast to implant failure, middle-ear effusion can be treated with myringotomy or may be allowed to resolve on its own. Finally, many ears that undergo ossiculoplasty are prone to the growth of soft-tissue adhesions or fibrosis. While revision surgery is possible in these ears, the long-term results of revision surgery are likely to be poor, since the adhesions or fibrosis will likely recur [18–19]. Were it known that soft-tissue fixation was the source of failure of middle-ear surgery, many clinicians would recommend alternative non-surgical treatments.

OCT middle ear imaging and vibrometry offer a potential solution to the diagnostic problem of differentiating these various conditions of the post-surgical ear. OCT vibrometry can be precisely targeted to structures of interest under OCT imaging guidance to make frequency-resolved measurements of their motion in response to sound. Generally, these measurements are reported as *mobility*, i.e., a point velocity normalized to ear canal pressure in units of mm/s/Pa [20]. The ability of OCT vibrometry to directly interrogate the mobility of middle ear structures offers benefits over tympanometry, which can only provide a measurement of the motion of the TM as a whole so that small changes in the mobility of individual structures like the ossicles are often overwhelmed by variability in the mechanical properties of the TM [21]. OCT vibrometry also offers advantages over laser Doppler vibrometry (LDV), a similar technology that has seen limited clinical use [22]. LDV has been shown to differentiate ossicular discontinuity from stapes and malleus fixation [23]. Its scanning and 3D extensions also enable full-field TM mobility mapping [24] and multidirectional vibrometric assessment of the target structures [25]. However, LDV still lacks co-registered, depth-resolved image guidance [26] and depth-resolved vibration measurements, both of which are achievable using OCT vibrometry.

The objective of the present study is to determine whether middle-ear OCT vibrometry can differentiate the causes of recurrent CHL after ossiculoplasty. Specifically, we aimed to: (1) establish condition-specific OCT vibrometry mobility signatures for effusion, soft-tissue fixation, and prosthesis disconnection at the umbo and at the center of the cartilage graft across six stimulus frequencies (500-3000 Hz); (2) quantify the magnitude of mobility for each condition as a function of frequency and test whether this feature can be used to diagnostically classify the condition of the ear; and (3) assess the implications of these signatures for clinical decision-making in the context of revision surgery.

## Materials and Methods

### Surgical Preparation

All cadaveric procedures in this study were performed under a protocol approved by the Dalhousie University Research Ethics Board (FILE #1034135). Fresh-frozen human cadaveric temporal bones were thawed and then prepared by removing the external soft tissues on the mastoid while preserving the ear canal, TM, and pinna. Cartilage was harvested from the tragus and/or concha of the pinna for later use as a cartilage graft. The ear canal, as well as the surface of the TM, was cleared of cerumen and other debris. Any liquid accumulated on the surface of the TM and around the annulus was removed by wicking with tissue paper. A cortical mastoidectomy with a wide posterior tympanotomy was performed to expose the middle ear through both the facial recess and the enlarged aditus to the mastoid antrum. The incudostapedial joint, incudomalleolar joint, and posterior incudal ligament were sectioned, followed by the removal of the buttress. The incus was then removed from the middle-ear space, leaving the stapes with its intact superstructure, intact malleus, and intact TM in place. A Kurz 2.5-mm Dresden clip partial ossicular replacement prosthesis (PORP) (Heinz Kurz GmbH, Dusslingen, Germany) was crimped onto the head of the stapes, and the space between the PORP plate and the posterosuperior quadrant of the TM was packed with cartilage harvested from the specimen’s concha or tragus, which was cut into a bespoke polygonal shape designed to cover the TM. The cartilage graft was shaved to a thickness appropriate to achieving simultaneous contact with the prosthesis and the TM with a degree of tension qualitatively representative of that achieved in typical surgical practice. Typically, this graft thickness was between 1.3 mm and 1.5 mm. During placement, the center of the PORP plate was visually aligned with the center of the cartilage graft (Fig. 1a). The near-planar, slightly convex surface of the cartilage graft was oriented laterally toward the TM, whereas the slightly concave surface was oriented medially toward the PORP plate, promoting conformal apposition at both interfaces. Figure 1b shows a photograph (lower inset) of the preparation, taken under a surgical microscope through the facial recess, and a schematic (upper inset) showing the placement of the prosthesis. Though not representative of typical clinical practice, the retrotympanic approach chosen for this model was important because it kept the TM intact, thereby reproducing the acoustics of a healed tympanoplasty better than could have been achieved with a transcanal approach involving tympanotomy.

**Fig. 1.**
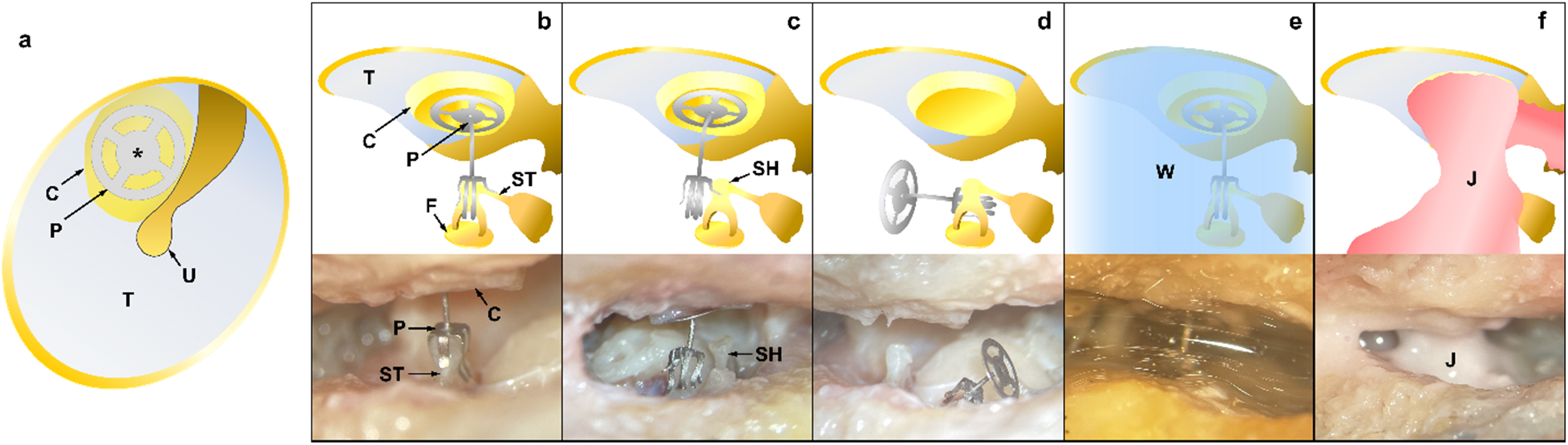
**(a)** *En-face* schematic view of the TM after PORP implantation. The cartilage graft lies beneath the TM, and the PORP beneath the cartilage graft. The cartilage graft is rendered partially transparent to show the PORP plate. The center of the PORP plate is visually aligned with that of the cartilage graft (asterisk). Schematics (upper row) and photos (lower row) taken through the facial recess showing examples of a middle ear prepared in **(b)** the baseline condition, **(c)** the PORP disconnected from the stapes, **(d)** the PORP disconnected from the TM, **(e)** the middle ear filled with water simulating effusion, and **(f)** the PORP surrounded with alginate gel (Jeltrate) simulating soft-tissue adhesion. C, cartilage graft; P, partial ossicular replacement prosthesis (PORP); T, tympanic membrane (TM); U, umbo; F, footplate of the stapes; ST, stapedius tendon; SH, stapes head; W, water; J, alginate gel (Jeltrate)

### OCT System

OCT measurements were made using a previously described [26] custom, handheld OCT system comprising an endoscopic handpiece and a swept-source OCT engine. The measurement protocols were designed for clinical translatability and are nearly identical to those used in ongoing clinical studies in patients. As shown in Fig. 2a, cadaveric temporal bones were imaged with intact pinnae and without making surgical modification to the external ear. Vibrometric displacement of the target structure (Fig. 2b) can be extracted from the acquired measurements, serving as the basis for subsequent mobility estimation. A standard, 4-mm otoscopic disposable plastic speculum (Heine: B-000.11.137., Heine Optotechnik, Gilching, Germany) was used to keep the geometric constraints and field of view the same as for clinical measurements. The handpiece contained a speaker and microphone to allow controlled delivery of sound and a black-and-white camera for *en*-*face* imaging of the TM. The OCT system was built around a swept-source laser (Insight Photonics SLE-101, Lafayette, Colorado) operating at a center wavelength of 1550 nm with a 40-nm sweep range at a 200-klines/s sweep rate. The system delivered an axial resolution in air of 37 μm, a lateral resolution in air of 40 μm, an axial dynamic range of 48 dB, and an axial depth range of 13 mm. A 2D micro-electro-mechanical systems (MEMS) scanning mirror (Mirrorcle, Richmond, California) located in the handpiece scanned the beam over a ±30° field of view in both lateral dimensions to form 2D and 3D OCT images. The handpiece was manually aligned in the ear canal so that the structure whose mobility was being measured was centered on the field of view in the camera image. The camera and OCT image were used to guide the operator in sequentially targeting two structures of interest: the umbo of the malleus and the center of the cartilage graft covering the prosthesis. The cartilage center was estimated from the shape of the cartilage graft as observed on the otoscopic camera image looking down the ear canal through the TM. For all measurements, the handpiece was manually held by the operator during alignment and measurement, simulating the intended use of the handpiece during clinical measurements.

**Fig. 2.**
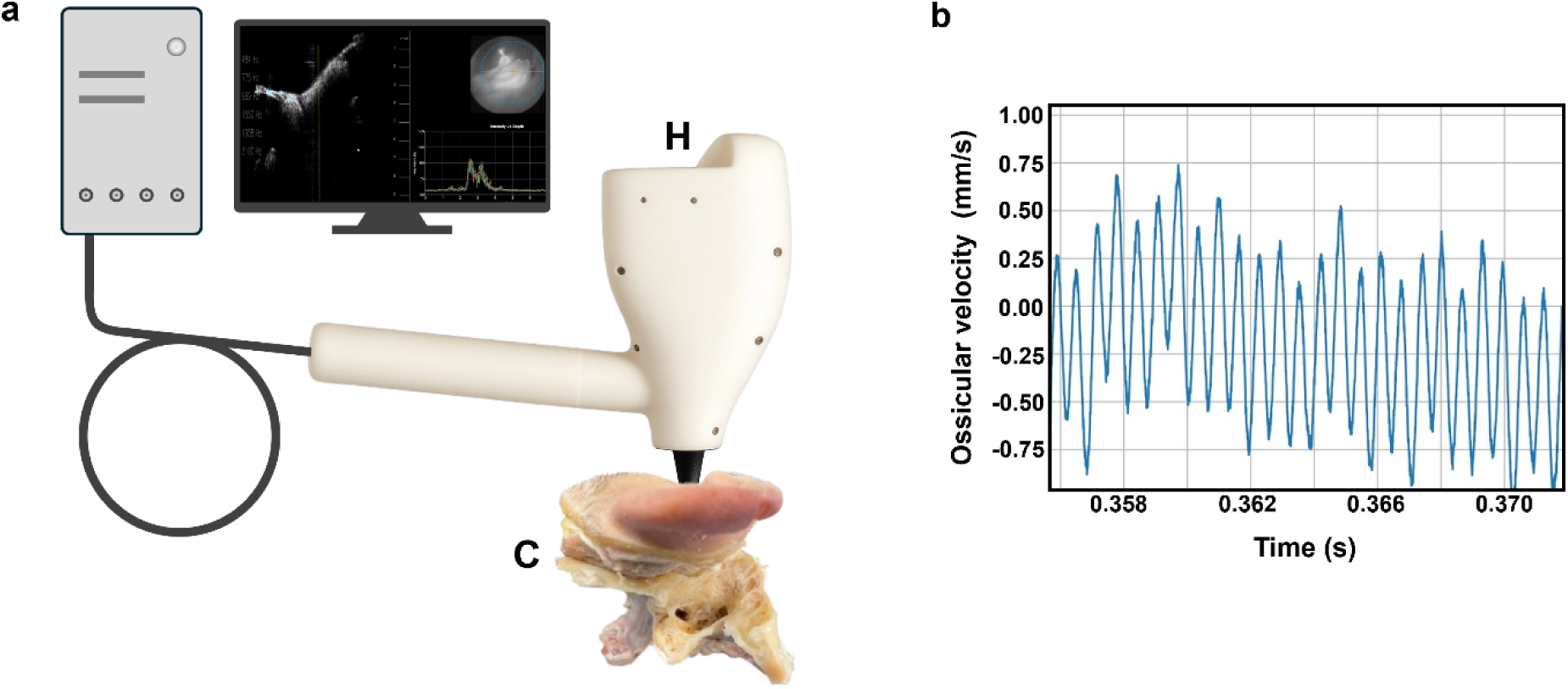
**(a)** Schematic of the OCT system imaging a cadaveric temporal bone. **(b)** Plot of the vibrometric displacement of the target structure extracted from the acquired measurements. H, OCT handpiece; C, cadaveric temporal bone

### Vibration Measurement

Vibrometric measurements in response to pure-tone sinusoidal stimuli were made at frequencies of 484 Hz, 775 Hz, 968 Hz, 1550 Hz, 1938 Hz, and 3100 Hz, selected as the closest frequencies to the standard audiometric frequencies 500 Hz, 750 Hz, 1000 Hz, 1500 Hz, 2000 Hz, and 3000 Hz that could be achieved given the frequency resolution of the sound generation system. At each frequency, the sound stimulus was presented for two seconds at 100 dB SPL, as measured by a microphone located at the tip of the handpiece. The system’s software user interface displayed video-rate B-mode images of the ear with a distal-proximal line drawn through the center of the B-mode image, indicating where depth-resolved vibrometric measurements would be performed. The operator would move the probe to align this line with the structure being vibrometrically interrogated and would then initiate a measurement by pressing a foot pedal. This would cause the B-mode image to be captured, mirror scanning to be stopped, and a sequence of OCT A-line measurements phase-locked with the sound stimulus to be obtained along a line intersecting the structure of interest. During the measurement, six pure-tone stimuli at the frequencies listed above would be presented sequentially into the ear canal through the handpiece speaker, while the OCT system simultaneously collected vibrometric responses. Following the measurement, the A-lines were automatically segmented using a peak-finding algorithm to select out the strongest reflector in the A-lines. At each pixel within the selected segment, the difference between the optical phases of the OCT signal measured in sequential A-lines was calculated. These phases were modulated by the acoustic displacement of the structure in response to the sound stimulus. The discrete Fourier transform (DFT) of this phase-difference signal was then calculated to obtain the amplitude of vibration at the stimulus frequency. From this DFT amplitude, the root-mean-squared (RMS) velocity magnitude of the pixel was calculated using the standard formula [4,6],

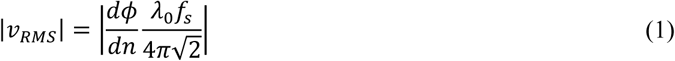

where 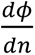 is the phase difference between subsequent A-lines (indexed by *n*) in radians at the target depth, *λ*_0_ = 1550 nm is the center wavelength of the laser sweep, and *f*_s_ = 200 kHz is the A-line rate of the swept source laser. Phase synchronization between the acoustic stimulus and laser sweeping was maintained throughout the measurement by generating the sound signal using a direct digital synthesizer clocked by the laser sweep clock [6]. In benchtop testing on a stationary target, this system was found to have an RMS vibrometric dynamic displacement noise floor of 255 pm for a two-second measurement.

The velocity magnitude at each pixel along the line through the structure of interest was then averaged over the 10 brightest pixels in the structure to obtain the final velocity estimate. This calculated velocity was not corrected for incidence angle, so the reported velocity should be interpreted as the component of the velocity along the optical beam axis rather than the velocity magnitude of the measured structure. Consequently, individual differences in the angle between the optical axis and the TM—which can range from 45**°** to 60**°**—will contribute about ±1.5 dB of inter-subject variation to the mobility measurements [27].

Supplemental Material 1 provides a system software video screen capture during a measurement sequence, showing the initial alignment of the system with the umbo, the completion of a measurement sequence, and the estimation of the mobility as a function of frequency.

Following measurement of the mobility at the umbo and at the center of the cartilage for each bone in its baseline implanted PORP condition, each bone was prepared in a sequence of simulated clinical conditions. The vibrometric measurements were repeated no less than three times at the same six frequencies at the umbo and cartilage locations in each condition. A 3D OCT image of the middle ear was also collected for each preparation. The bone was periodically moistened during measurements to prevent dehydration.

The following post-surgical failure conditions were simulated. First, to simulate a disconnection of the prosthesis, the base of the PORP was disconnected from the stapes head, while the plate remained attached to the cartilage beneath the TM by surface tension. Second, the PORP plate was also disconnected from the cartilage and left prone in the middle-ear cavity to simulate a complete disconnection of the PORP from the TM. Third, the PORP was re-mounted onto the stapes head and re-attached to the cartilage to restore it to its baseline condition. A new baseline measurement was taken to confirm that middle-ear mechanics had returned to the baseline level. One-way ANOVA found no significant difference between the initial baseline and subsequent baseline measurements (*p* > 0.05) across frequencies and locations. After that, water was injected into the middle-ear cavity from the facial recess to fill the middle ear cavity with fluid, simulating middle-ear effusion, and mobility measurements were taken. Fourth, the water was suctioned and wicked from the middle-ear cavity, and a third baseline measurement was performed. Following this third baseline, alginate gel (Jeltrate, Dentsply Caulk, Milford, Delaware) was mixed to produce a consistency qualitatively similar to middle-ear mucosa. The alginate gel was placed around the PORP and the remaining ossicles (i.e., malleus and stapes) to simulate soft-tissue fixation. This alginate-gel model of soft-tissue fixation has been used in previous studies of middle-ear mobility in states of pathology [28]. Schematics and photos showing these four conditions can be seen in Fig. 1c-f.

Each of these four conditions was modeled in a minimum of *N* = 10 temporal bones, although not all bones were prepared in all the conditions due to incidental damage (e.g., TM perforation) that occurred to some of the bones during preparation of the various middle ear conditions and which excluded them from further measurements.

### Data Analysis

For each of these conditions, a total of 12 “features” were measured, namely the mobility measurements taken at two locations (umbo and center of cartilage graft) and at six frequencies. These features were used to train a Random Forest classification model to distinguish the conditions (simulated effusion, simulated soft-tissue fixation, and disconnection), and the model was then used to predict the condition of the middle ear from the features, both when all of the data were used to train the model and in a leave-one-out cross-validation paradigm. Although two distinct disconnection conditions (disconnected at the stapes only and disconnected at the TM) were created, an ANOVA test found no significant difference in their mobilities (*p* > 0.05 for all frequencies), and so they were pooled into a single condition of “Disconnection” in further analysis.

In leave-one-out cross-validation, all but one of the datasets are used to train the model, and then the model is used to predict the condition of the dataset that it was not trained on from the features. This is repeated with each dataset being the one left out. Then correct and incorrect results are tallied to produce an overall accuracy score. Data were also aggregated for each pathological condition (i.e., effusion, soft-tissue fixation, disconnection at the stapes, and disconnection at the TM) and compared against the baseline conditions. The baseline condition at the start of the experiment and after simulating and then reversing the effusion and soft-tissue fixation conditions were compared to each other using one-way ANOVA at each frequency and location, and no statistically significant differences between them were observed at a *p* < 0.05 level for any frequency. Because of the lack of significant differences, the baseline measurements were pooled in the model training data.

## Results

The statistics of mobility measurements from all the samples in both baseline (aggregated from all three baseline measurements) and simulated conditions are detailed in Table 1. Compared to the pooled baseline mobility values, the conditions simulating soft-tissue fixation and effusion both led to a broadband mobility decrease of 8.0 dB and 17.5 dB, respectively, when averaged across six stimulus frequencies and two measurement locations. The conditions simulating prosthesis disconnections at the stapes and the TM, by contrast, led to a low-frequency (at 500 Hz and 750 Hz) mobility increase of 9.9 dB and 11.1 dB, respectively, when averaged across these two frequencies and two measurement locations.

**Table 1.** Descriptive statistics for the experiment. Numbers represent the mean value, median value, and standard deviation of the mobilities at every particular location and every individual frequency. The mean value, median value, and standard deviation of mobility are calculated in dB relative to 1 mm/s/Pa. Baseline data were pooled across the initial baseline, the baseline obtained before simulating effusion, and the baseline obtained before simulating fixation. Sample size information is provided in the bottom row

| Stimulus frequency |  | Baseline (Pooled) |  | Simulated fixation |  | Simulated effusion |  | Disconnected at stapes |  | Disconnected at TM |  |
| --- | --- | --- | --- | --- | --- | --- | --- | --- | --- | --- | --- |
|  |  | Umbo | Cartilage graft | Umbo | Cartilage graft | Umbo | Cartilage graft | Umbo | Cartilage graft | Umbo | Cartilage graft |
| 484 Hz | Mean | -17.9 | -24.5 | -30.1 | -33.2 | -35.9 | -37.3 | -6.5 | -4.5 | -4.3 | -1.6 |
|  | Median | -17.2 | -24.3 | -31.1 | -37.2 | -36.0 | -38.2 | -5.7 | -4.0 | -3.7 | -2.6 |
|  | Std. | 5.3 | 5.3 | 7.3 | 8.6 | 3.0 | 4.0 | 4.9 | 4.3 | 5.8 | 5.8 |
| 775 Hz | Mean | -10.7 | -19.9 | -24.8 | -26.2 | -35.7 | -35.8 | -10.8 | -11.7 | -11.9 | -11.0 |
|  | Median | -10.3 | -19.7 | -26.8 | -30.4 | -35.7 | -35.9 | -10.0 | -12.1 | -12.0 | -9.3 |
|  | Std. | 5.9 | 5.6 | 6.7 | 11.0 | 2.8 | 4.8 | 5.2 | 3.7 | 5.7 | 5.3 |
| 968 Hz | Mean | -10.0 | -17.6 | -21.7 | -25.9 | -34.2 | -33.2 | -10.7 | -15.0 | -11.5 | -14.9 |
|  | Median | -8.9 | -18.9 | -24.0 | -29.3 | -34.4 | -35.4 | -10.1 | -14.7 | -10.8 | -13.5 |
|  | Std. | 4.9 | 5.7 | 7.8 | 8.4 | 4.6 | 5.9 | 5.2 | 3.4 | 5.2 | 3.6 |
| 1550 Hz | Mean | -9.6 | -15.9 | -18.5 | -24.5 | -32.1 | -30.2 | -11.1 | -15.9 | -10.5 | -15.7 |
|  | Median | -10.0 | -15.7 | -18.8 | -25.5 | -32.4 | -32.1 | -12.4 | -16.3 | -11.5 | -16.1 |
|  | Std. | 4.1 | 4.9 | 5.8 | 6.1 | 6.6 | 6.1 | 6.4 | 4.0 | 6.2 | 4.5 |
| 1938 Hz | Mean | -11.6 | -18.3 | -17.4 | -23.2 | -35.4 | -32.7 | -11.5 | -17.8 | -11.2 | -16.9 |
|  | Median | -13.3 | -18.3 | -17.5 | -25.2 | -36.4 | -33.3 | -12.3 | -17.7 | -12.0 | -17.8 |
|  | Std. | 6.6 | 6.0 | 4.4 | 6.0 | 4.9 | 3.3 | 5.5 | 4.4 | 6.2 | 3.9 |
| 3100 Hz | Mean | -10.4 | -19.1 | -12.2 | -23.9 | -26.3 | -26.5 | -8.3 | -21.5 | -6.9 | -19.9 |
|  | Median | -9.0 | -19.1 | -15.4 | -25.5 | -25.7 | -27.2 | -6.9 | -22.4 | -5.0 | -20.7 |
|  | Std. | 9.0 | 9.4 | 8.3 | 6.2 | 6.8 | 6.4 | 7.5 | 7.0 | 8.7 | 7.2 |
| Sample Size |  | N = 32 |  | N = 10 |  | N = 10 |  | N = 13 |  | N = 11 |  |

All mobilities are expressed in decibels (dB) relative to 1 mm/s/Pa.

Figure 3 shows measured mobilities at baseline (with the PORP installed and connected to the TM and the stapes) and in the various simulated post-surgical disorders. Significant decreases from baseline were observed at all frequencies for the simulated effusion condition at the umbo (*p* < 0.01 at 3000 Hz; *p* < 0.001 at the other five frequencies) and at all frequencies except 3000 Hz at the cartilage graft (*p* < 0.01 at 1500 Hz; *p* < 0.001 at the other four frequencies). For the simulated soft-tissue fixation, significant decreases from baseline were observed at 500 Hz, 750 Hz, 1000 Hz, and 1500 Hz at the umbo (*p* < 0.01 for these four frequencies) and at 500 Hz, 1000 Hz, and 1500 Hz at the cartilage graft (*p* < 0.05 for these three frequencies). The disconnections at the stapes and the TM were found not to be statistically different from each other and so were pooled together in further analysis, including classification analysis. However, the conditions of disconnected at the stapes and disconnected at the TM each individually showed significantly higher mobility than baseline at 500 Hz at the umbo (*p* < 0.001 for both conditions) and at 500 Hz (*p* < 0.0001 for both conditions) and 750 Hz (*p* < 0.001 for disconnected at the stapes and *p* < 0.01 for disconnected at the TM) at the cartilage graft.

**Fig. 3.**
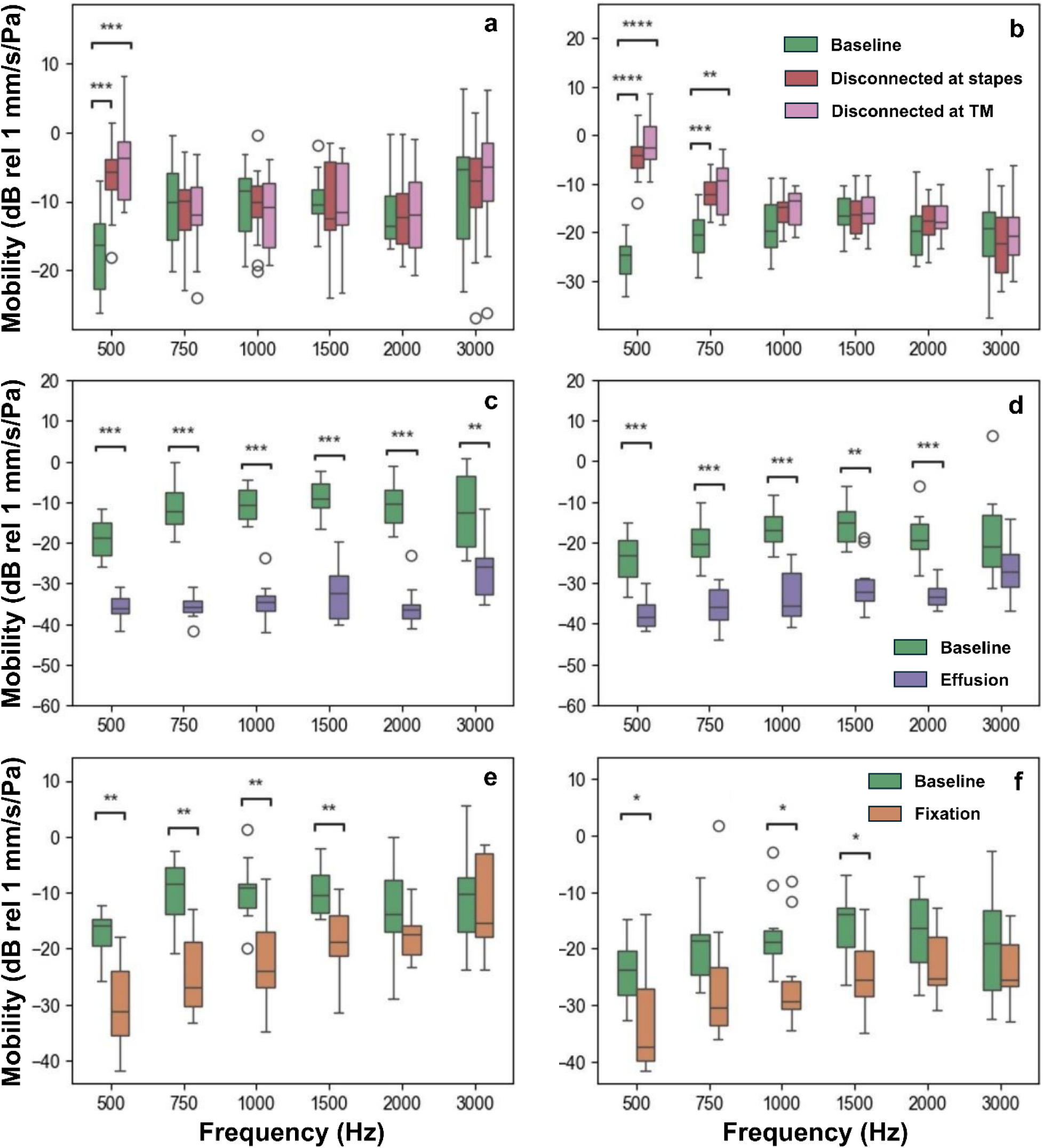
Measured mobilities of temporal bones prepared in baseline (PORP implanted with good coupling to the TM and stapes), PORP disconnected at the stapes, PORP disconnected at the TM, shown in **(a)** and **(b)**; simulated effusion achieved by filling the ear with water, shown in **(c)** and **(d)**; and simulated soft-tissue fixation achieved with alginate gel, shown in **(e)** and **(f)**. Significant differences as determined by a Mann-Whitney test are denoted by **** (*p* < 0.0001), *** (*p* < 0.001), ** (*p* < 0.01), and * (*p* < 0.05). The insets **(a)**, **(c)**, and **(e)** show the results at the location of the umbo, while the insets **(b)**, **(d)**, and **(f)** show the results at the location of the cartilage graft

Figure 4 shows the change in mobility from baseline for each of the conditions. The mobility change from baseline is a potentially more sensitive diagnostic measure as it compares the measurements under each condition with its corresponding baseline within the same bone, rather than with the baseline measurements across different bones. It also provides a model for a clinical use case in which degraded hearing can be interrogated by comparing it against baseline OCT vibrometry data collected after surgery.

**Fig. 4.**
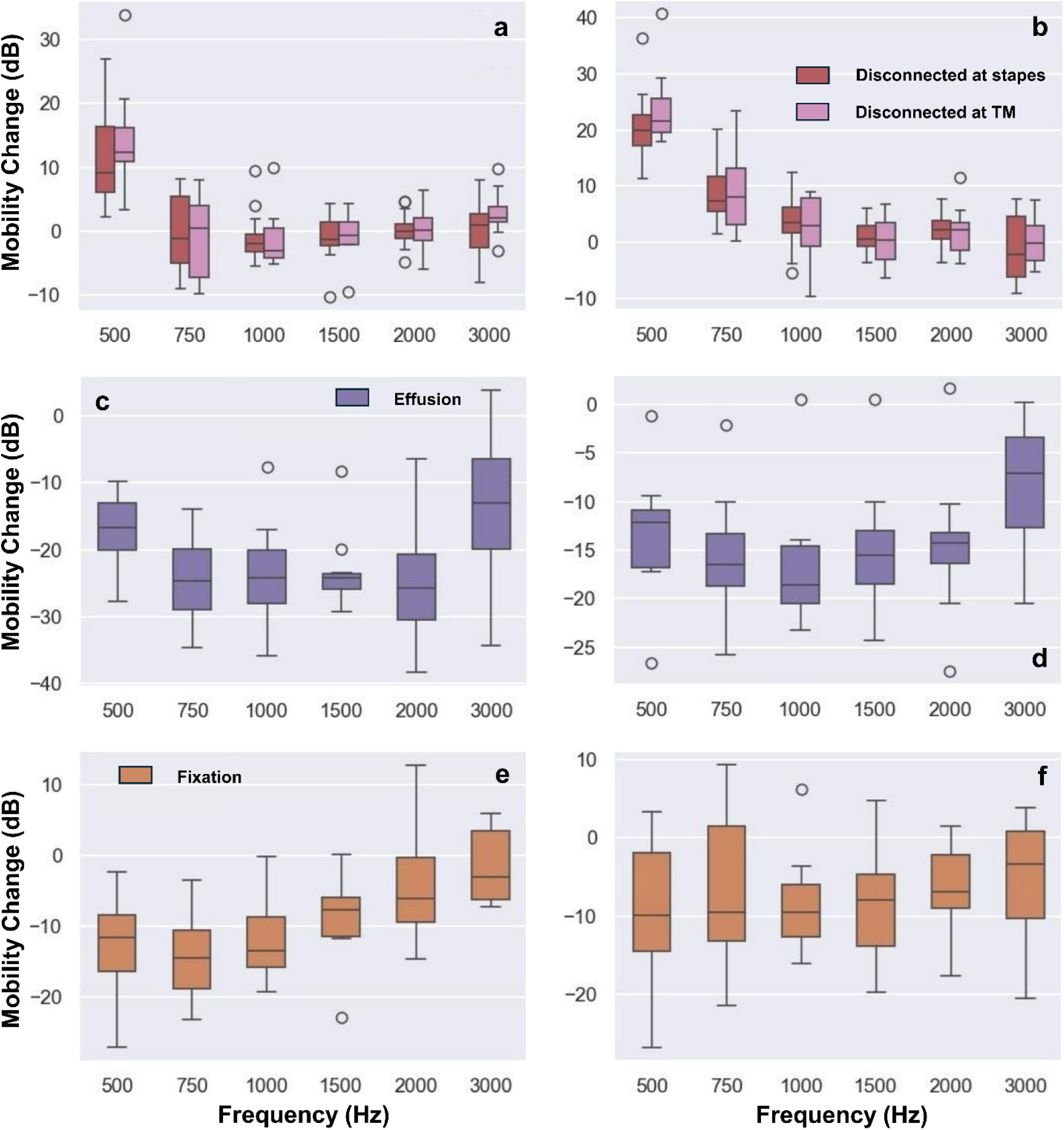
Changes in mobility relative to baseline condition for the simulated post-surgical conditions: **(a)** PORP disconnected at the stapes and at the TM, measured at the umbo; **(b)** PORP disconnected at the stapes and at the TM, measured at the cartilage graft; **(c)** effusion, measured at the umbo; **(d)** effusion, measured at the cartilage graft; **(e)** soft-tissue fixation, measured at the umbo; and **(f)** soft-tissue fixation, measured at the cartilage graft

Multiple linear regression analysis showed that there was a significant difference in the slope of the mobility-versus-frequency curve between the simulated soft-tissue fixation and the simulated effusion at the umbo (*p* < 0.0001) but not at the cartilage graft. This finding indicates that the frequency-independent reduction in mobility observed for effusion (Fig. 4c) is distinguishable from a mobility that increases with increasing frequency for simulated soft-tissue fixation (Fig. 4e).

Figure 5 shows the results of Random Forest classification, in which a Random Forest classifier was trained to classify the bone condition from the mobility measurements. Figure 5a shows the result of applying a linear discriminant analysis to project data from the 12-dimensional feature space generated by the mobility measurements (six frequencies × two locations) onto a reduced 2D space. A Random Forest classifier was then trained on the full dataset and applied to classify the disorders. All bone conditions are correctly classified (accuracy = 100%), which shows that high classification accuracy is possible, although training the model using all available data introduces the possibility of overfitting and may cause the accuracy to exceed the accuracy that would be seen for new samples that were not part of the training set. One pair of samples lies very close to the border between simulated effusion and simulated soft-tissue fixation, indicating that their correct assignment was likely a case of overfitting. To assess the ability of the model to predict the state of middle ears that were not in the training set, we performed leave-one-out cross-validation, in which the model was trained on all but one of the datasets and then asked to predict the condition of the left-out dataset from the mobilities. This was repeated with each dataset being left out once, and the total accuracy of the predictions tabulated. The accuracy obtained from this leave-one-out cross-validation test was typically 90.9% (40/44 datasets were correctly predicted), although a variation of the accuracy ranging from 88.6% (39/44) to 93.2% (41/44) was observed across data-analysis program runs due to the stochastic nature of the random forest model [29]. Even with the same total accuracy, the accuracy for each individual condition might be different. The condition simulating effusion typically showed the lowest accuracy, while the disconnection condition typically showed the highest. Two representative confusion matrices from cross-validation showing the same total accuracy (90.9%) but different individual-condition accuracies can be found in Fig. 5b.

**Fig. 5.**
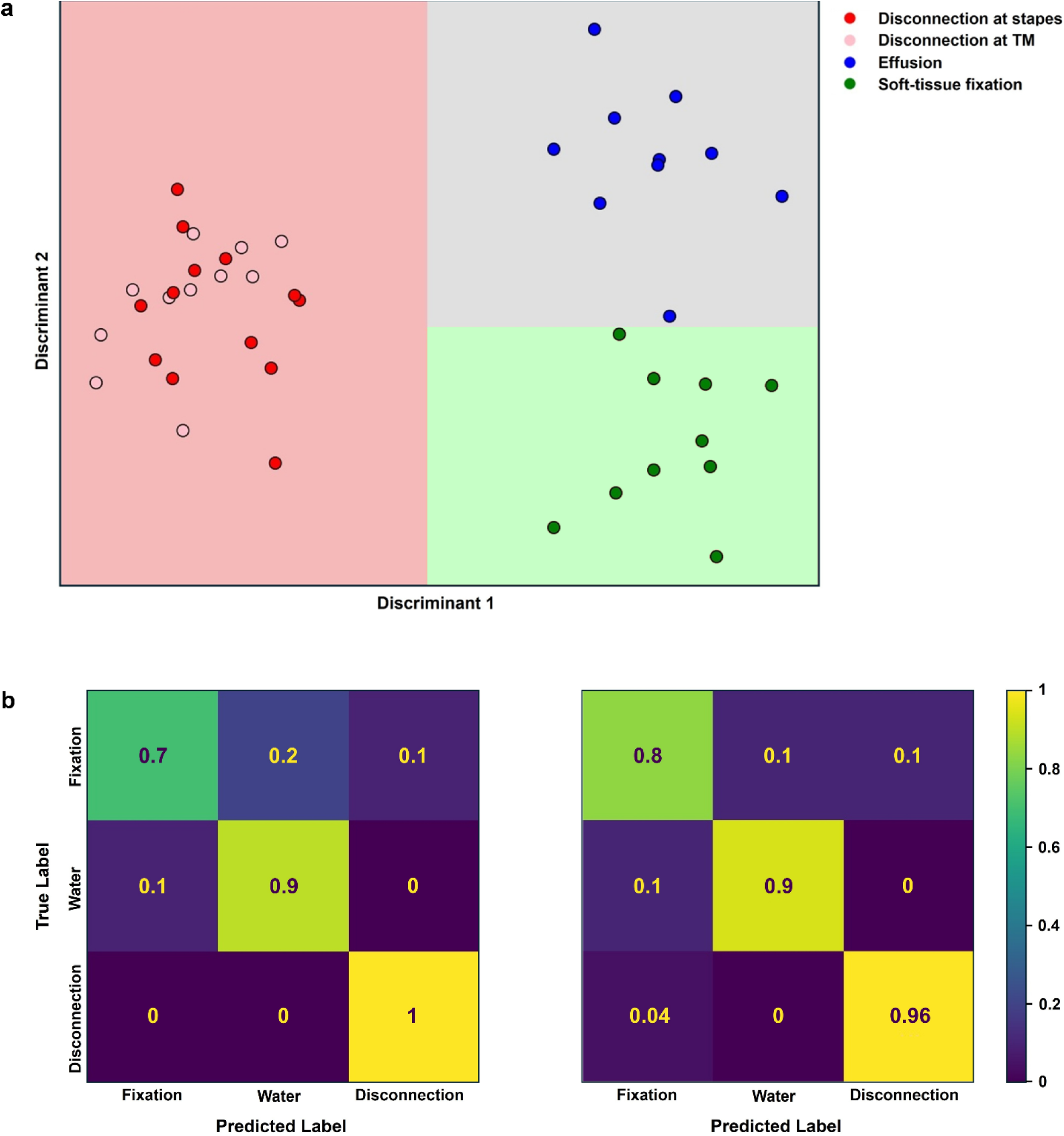
Random Forest classification results for predicting middle-ear conditions from mobility measurements: **(a)** classification in 2D space using linear discriminant analysis; **(b)** representative confusion matrices from leave-one-out cross-validation of a Random Forest classifier trained on all 12 features

The relative feature importance was assessed using the Gini Importance, which calculates the average model fit accuracy decrease from removing a feature from the input. Again, because of the stochastic nature of the random forest method, the distribution of Gini Importance among features varied across data-analysis program runs. Figure 6 shows representative Gini Importance values for each combination of frequency and location. Consistent with Figs. 3 and 4, low-frequency features, particularly the 500-Hz mobility at the umbo and cartilage graft, appear most important for classifying the middle ear state from mobility data.

**Fig. 6.**
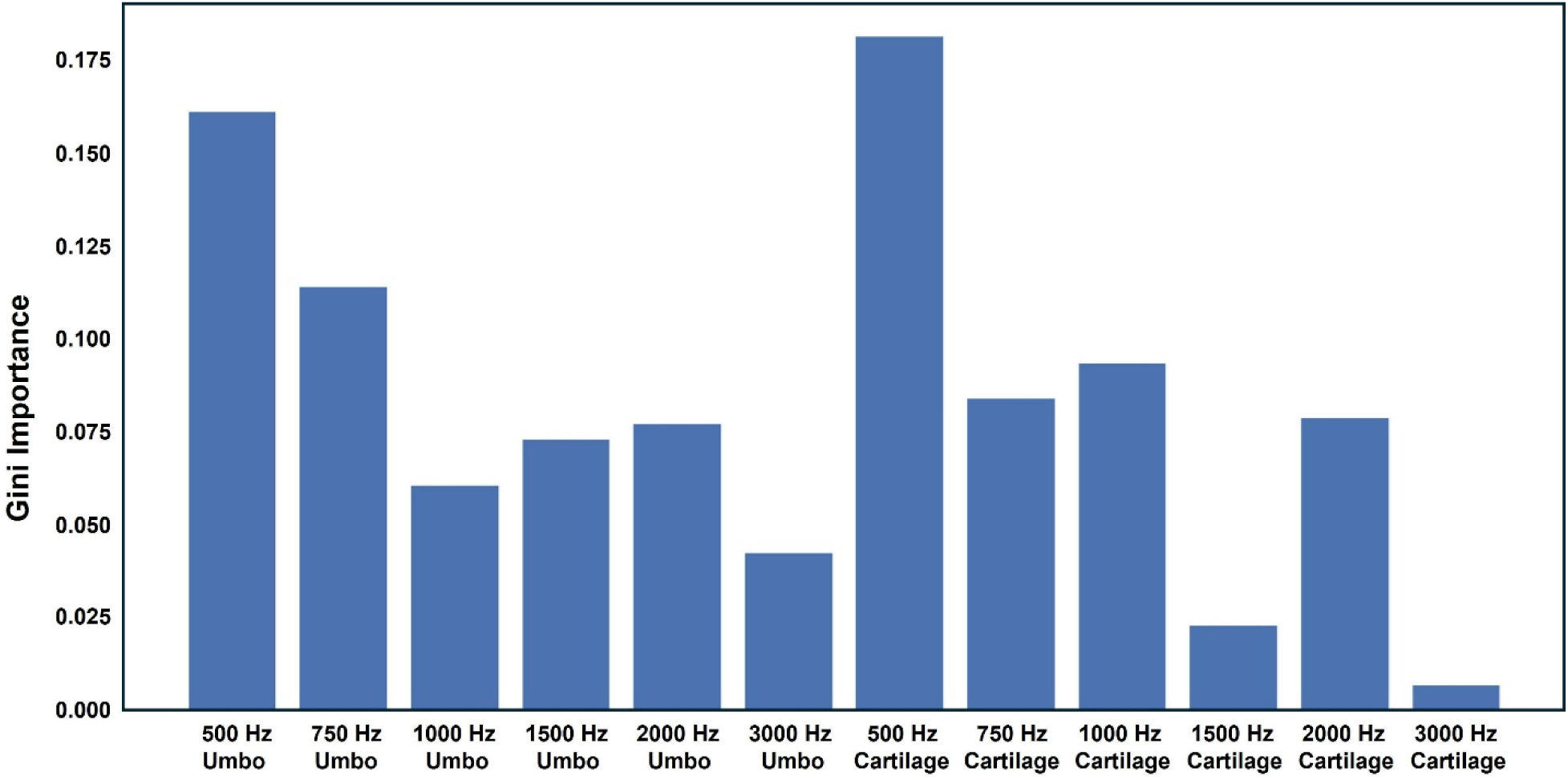
Representative Gini importance values across the 12 features used in Random Forest classification

## Discussion

The results of the experiment show that various causes of recurring CHL in a post-surgical ear can be accurately differentiated using OCT vibrometry. Aggregated data show significant decreases in the mobility from simulated effusion and soft-tissue fixation as one would expect both on general mechanical principles and from previous laser Doppler vibrometry measurements that have studied the effects of simulated otitis media [30] and simulated soft-tissue fixation using alginate gel [28], although these experiments are not directly comparable because they looked at ears with intact ossicular chains rather than a simulated post-operative ear with a prosthesis. The result that disconnection causes a low-frequency increase in mobility also makes mechanical sense. Dislocation of the PORP results in loss of coupling between the TM and the stapes. With the TM then coupled only to the underlying cartilage graft and the isolated malleus, the load on the TM is reduced compared with the properly positioned PORP, both of which couple the TM to the inner ear through the stapes. This reduction in loading explains the observed increase in low-frequency mobility. To our knowledge, there are no previously published studies examining the vibrometric effects of ossicular prosthesis disconnection, although one of our co-authors previously investigated the impact of progressive fibrotic fixation on ossicular mobility using LDV [31].

While these differences in the aggregated data are suggestive of diagnostic utility, the analysis of the predictability of middle-ear conditions from mobilities in a Random Forest classification model demonstrates that the middle-ear status can be determined with high reliability using OCT vibrometry in individual ears, at least for the cadaveric model used in this study. However, leave-one-out cross-validation indicates that discriminating simulated effusion from simulated soft-tissue fixation is more difficult than distinguishing either from prosthesis disconnection. This is an intuitive result since prosthesis disconnection causes an increase in mobility while both fixation and effusion cause a decrease in mobility.

This study confined itself to measurements performed in a cadaveric temporal bone model of the post-surgical ear and, as a result, some important differences between the preclinical and clinical situations should be noted.

First, we used a retrotympanic approach to insert the implant and used tragal or conchal cartilage to secure the implant plate against the TM. Clinically, most middle-ear surgeries involving PORP implantation are performed with a transcanal approach with an underlay or palisade tympanoplasty, depending on circumstances and surgeon preference [32]. We opted for the retrotympanic approach to preserve the integrity of the TM and to reproduce the acoustics of a middle ear sealed off from the ear canal by a healed TM. However, this choice meant that we were inserting the prosthesis and cartilage against the TM under tension rather than under a flaccid tympanomeatal flap as would be the case in a typical transcanal surgery. We believe that our retrotympanic implantation reproduces the acoustics of a healed ear sufficiently well to support our conclusions because the TM tension will return after clinical transcanal tympanoplasty as the TM heals and integrates the cartilage graft. Post-surgical healing has other effects that were not captured by our cadaveric model. In a healed post-surgical ear, the cartilage graft is integrated into the TM, often with various degrees of fibrous tissue that may affect the mobility of the graft. Scarring and stiffening also occur within the middle-ear joints and around the implant and can affect implant and TM mobility. These effects were not captured in our model.

In modeling effusion, we filled the middle ear with water. Water is a reasonable model for the effusion in serous otitis media, but not for acute or adhesive otitis media, where effusion is more viscous [13]. However, these more viscous effusions could be expected to lead to an even greater decrease in mobility than the condition tested in this study and so should be at least as detectable with OCT vibrometry as our simulated effusion.

Alginate has previously been used in cadaveric models to simulate soft-tissue fixation [28]. While the mechanical properties of alginate qualitatively resemble those of middle-ear adhesions, alginate’s properties depend strongly on the details of its preparation, particularly the time it is allowed to cure, and we did not attempt to quantitatively control either consistency or mechanical similarity to middle-ear mucosa, relying instead on a qualitative assessment of the alginate consistency. So, while the finding that fixation with alginate resulted in a mobility that rose from low to high frequency, which could be distinguished from the frequency-independent mobility decrease from the simulated effusion, it is not clear whether similar behavior will be found in the clinic. The present study focused on postoperative scenarios most relevant to a PORP reconstruction model, where fibrous fixation, rather than bony fixation, is expected because middle-ear prostheses tend to induce tissue fibrosis after implantation [33]. In contrast, bony fixation is typically associated with pre-operative middle-ear diseases such as otosclerosis and would not be expected to occur following a PORP reconstruction [34].

In simulating the disconnection of the PORP from the stapes, we avoided contact between the PORP clip and any surrounding structures. In clinical cases, the PORP can sometimes fail in a way that leaves the clip in contact with the promontory, and this may create a stiffer-than-normal PORP and a decrease in PORP mobility rather than an increase as was seen in the present experiment. This scenario was not simulated in the present study.

In this study, we employed a Random Forest classifier to demonstrate the ability of the vibrometric measurements to distinguish the state of the middle ear in cases of CHL recurrence following ossiculoplasty. While this general classification approach is suitable for clinical translation, because of the differences between the cadaveric temporal-bone model we used and the healed post-surgical ear, it is not clear whether the classification model trained on our cadaver data would accurately classify patient ears. Moreover, while for the purpose of this experiment, we were able to generate a set of ears with uniform preparation, the types of ears encountered in the clinic have a wide spectrum of degrees of fixation and prosthesis disconnection. This variation will likely make it more difficult both to train a classifier using patient data and to apply categorical labels to patients. Categorization of patients is also made difficult because confirmatory diagnosis of the patient’s condition may come only after revision surgery (for prosthesis disconnections and soft-tissue fixation), following expression of fluid via myringotomy (for effusion), or may not come at all (if the patient decides not to proceed to revision surgery). As a result, acquiring a suitable clinical dataset for model training may be challenging in practice.

Another state-of-the-art noninvasive approach for assessing middle-ear mechanics is wideband acoustic immittance (WAI). WAI quantifies ear-canal absorbance across frequency and is sensitive to middle-ear effusion [35], ossicular discontinuity, and stapes fixation [22]. Postoperative studies have also shown that WAI absorbance patterns change after tympanoplasty [36] or cartilage-graft repair [37], but to our knowledge, prosthesis-specific (e.g., PORP) patterns have not yet been characterized, nor has WAI been used to differentiate causes of post-surgical CHL.

By contrast, OCT vibrometry provides distinct signatures for various causes of post-surgical CHL, as demonstrated in this study. Disconnections can be identified by low-frequency hypermobility at the implant site; effusion by wideband attenuation at both the implant site and the umbo; and soft-tissue fixation by an attenuated mobility that increases with frequency. The overall trends observed here are consistent with middle-ear mechanical intuition and are therefore expected to hold in clinical cases as well, suggesting that OCT vibrometry offers a promising new direction for guiding decision-making following a poor outcome from middle-ear surgery.

## Statements and Declarations

## Ethical approval and informed consent

All human cadaveric temporal bones were used under a protocol approved by the Dalhousie University Research Ethics Board (FILE #1034135).

## Acknowledgements and funding

This study was funded by the Canadian Institutes of Health Research (CIHR), Project Grant #PJT180435.

## Notes

### Competing Interest Statement

Xiaojie Yang declares no conflicts of interest related to this work.
David Morris and Robert Adamson declare a financial conflict of interest related to their relationship with Carl Zeiss Meditec, with whom they are developing optical coherence tomography technology for the diagnosis of middle-ear disorders. Both David Morris and Robert Adamson are financially compensated for their work with Carl Zeiss.
No financial, personal, or professional relationships have influenced the design, conduct, or reporting of the research presented in this manuscript. All submitted materials, including the manuscript, figures, table, and supplementary video, are not associated with any conflicts of interest.

## References

[1] Tan HEI, Maria PLS, Wijesinghe P, Kennedy BF, Allardyce BJ, Eikelboom RH, Atlas MD, Dilley RJ (2018) Optical coherence tomography of the tympanic membrane and middle ear: A review. Otolaryngol Head Neck Surg 159(3): 424–438. 10.1177/0194599818775711.

[2] Cho NH, Lee SH, Jung W, Jang JH, Kim J (2015) Optical coherence tomography for the diagnosis and evaluation of human otitis media. J Korean Med Sci 30(3): 328–335. 10.3346/jkms.2015.30.3.328.

[3] Djalilian HR, Ridgway J, Tam M, Sepehr A, Chen Z, Wong BJF (2008) Imaging the human tympanic membrane using optical coherence tomography in vivo. Otol Neurotol 29(8): 1091–1094. 10.1097/MAO.0b013e31818a08ce.

[4] Kirsten L, Schindler M, Morgenstern J, Erkkila MT, Golde J, Walther J, Rottmann P, Kemper M, Bornitz M, Neudert M, Zahnert T, Koch E (2018) Endoscopic optical coherence tomography with wide field-of-view for the morphological and functional assessment of the human tympanic membrane. J Biomed Opt 24(3): art.031017. 10.1117/1.JBO.24.3.031017.

[5] Pitris C, Saunders KT, Fujimoto JG, Brezinski ME (2001) High-resolution imaging of the middle ear with optical coherence tomography - A feasibility study. Arch Otolaryngol Head Neck Surg 127(6): 637–642. 10.1001/archotol.127.6.637.

[6] MacDougall D, Farrell J, Brown J, Bance M, Adamson R (2016) Long-range, wide-field swept-source optical coherence tomography with GPU accelerated digital lock-in Doppler vibrography for real-time, in vivo middle ear diagnostics. Biomed Opt Express 7(11): 4621–4635. 10.1364/BOE.7.004621.

[7] Kim W, Kim S, Huang S, Oghalai JS, Applegate BE (2019) Picometer scale vibrometry in the human middle ear using a surgical microscope based optical coherence tomography and vibrometry system. Biomed Opt Express 10(9): 4395–4410. 10.1364/BOE.10.004395.

[8] Park J, Carbajal EF, Chen X, Oghalai JS, Applegate BE (2014) Phase-sensitive optical coherence tomography using an Vernier-tuned distributed Bragg reflector swept laser in the mouse middle ear. Opt Lett 39(21): 6233–6236. 10.1364/OL.39.006233.

[9] Van der Jeught S, Dirckx JJJ, Aerts JRM, Bradu A, Podoleanu AG, Buytaert JAN (2013) Full-field thickness distribution of human tympanic membrane obtained with optical coherence tomography. J Assoc Res Otolaryngol 14(4): 483–494. 10.1007/s10162-013-0394-z.

[10] Djalilian HR, Rubinstein M, Wu EC, Naemi K, Zardouz S, Karimi K, Wong BJF (2010) Optical coherence tomography of cholesteatoma. Otol Neurotol 31(6): 932–935. 10.1097/MAO.0b013e3181e711b8.

[11] Wang J, Chawdhary G, Yang X, Morin F, Khalid-Raja M, Farrell J, MacDougall D, Chen F, Morris D, Adamson RBA (2022) Optical clearing agents for optical imaging through cartilage tympanoplasties: A preclinical feasibility study. Otol Neurotol 43(4): e467–e474. 10.1097/MAO.0000000000003502.

[12] Wang J, Couvreur F, Farrell JD, Ghedia R, Shoman N, Morris DP, Adamson RBA (2025) Fusion of middle ear optical coherence tomography and computed tomography. JAMA Otolaryngol Head Neck Surg 151(5): 476–484. 10.1001/jamaoto.2025.0043.

[13] Monroy GL, Pande P, Shelton RL, Nolan RM, Spillman DR Jr, Porter RG, Novak MA, Boppart SA (2017) Non-invasive optical assessment of viscosity of middle ear effusion in otitis media. J Biophotonics 10: 394–403. 10.1002/jbio.201500313.

[14] Monroy GL, Shelton RL, Nolan RM, Nguyen CT, Novak MA, Hill MC, McCormick DT, Boppart SA (2015) Noninvasive depth-resolved optical measurements of the tympanic membrane and middle ear for differentiating otitis media. Laryngoscope 125(8): e276–e282. 10.1002/lary.25141.

[15] Wang J, Chawdhary G, Farrell J, Yang X, Farrell M, MacDougall D, Trudel M, Shoman N, Morris DP, Adamson RBA (2022) Transtympanic visualization of cochlear implant placement with optical coherence tomography: A pilot study. Otol Neurotol 43(8): e824–e828. 10.1097/MAO.0000000000003635.

[16] Cox MD, Page JC, Trinidade A, Dornhoffer JL (2017) Long-term complications and surgical failures after ossiculoplasty. Otol Neurotol 38(10): 1450–1455. 10.1097/MAO.0000000000001572.

[17] Cox MD, Trinidade A, Russell JS, Dornhoffer JL (2017) Long-term hearing results after ossiculoplasty. Otol Neurotol 38(4): 510–515. 10.1097/MAO.0000000000001339.

[18] Dornhoffer JL, Gardner E (2001) Prognostic factors in ossiculoplasty: A statistical staging system. Otol Neurotol 22(3): 299–304. 10.1097/00129492-200105000-00005.

[19] Yung M (2006) Long-term results of ossiculoplasty: Reasons for surgical failure. Otol Neurotol 27(1): 20–26. 10.1097/01.mao.0000176173.94764.f5.

[20] Whittemore KR, Merchant SN, Poon BB, Rosowski JJ (2004) A normative study of tympanic membrane motion in humans using a laser Doppler vibrometer (LDV). Hear Res 187(1-2): 85–104. 10.1016/s0378-5955(03)00332-0.

[21] Voss SE, Allen JB (1994) Measurement of acoustic impedance and reflectance in the human ear canal. J Acoust Soc Am 95(1): 372–384. 10.1121/1.408329.

[22] Nakajima HH, Pisano DV, Roosli C, Hamade MA, Merchant GR, Mahfoud L, Halpin C, Rosowski JJ, Merchant SN (2012) Comparison of ear-canal reflectance and umbo velocity in patients with conductive hearing loss: A preliminary study. Ear Hear 33(1): 35–43. 10.1097/AUD.0b013e31822ccba0.

[23] Rosowski JJ, Nakajima HH, and Merchant SN (2008) Clinical utility of laser-Doppler vibrometer measurements in live normal and pathologic human ears. Ear Hear 29(1): 3–19. 10.1097/AUD.0b013e31815d63a5.

[24] Zhang X, Guan X, Nakmali D, Palan V, Pineda M, and Gan RZ (2014) Experimental and modeling study of human tympanic membrane motion in the presence of middle ear liquid. J Assoc Res Otolaryngol 15(6): 867–881. 10.1007/s10162-014-0482-8.

[25] Dobrev I and Sim JH (2018) Magnitude and phase of three-dimensional (3D) velocity vector: Application to measurement of cochlear promontory motion during bone conduction sound transmission. Hear Res 364: 96–103. 10.1016/j.heares.2018.03.022.

[26] Farrell JD, Wang J, MacDougall D, Yang X, Brewer K, Couvreur F, Shoman N, Morris DP, Adamson RBA (2023) Geometrically accurate real-time volumetric visualization of the middle ear using optical coherence tomography. Biomed Opt Express 14(7): 3152–3171. 10.1364/BOE.488845.

[27] Fay JP, Puria S, and Steele CR (2006) The discordant eardrum. Proc Natl Acad Sci U S A 103(52): 19743–19748. 10.1073/pnas.0603898104.

[28] Nakajima HH, Ravicz ME, Merchant SN, Peake WT, Rosowski JJ (2005) Experimental ossicular fixations and the middle ear’s response to sound: Evidence for a flexible ossicular chain. Hear Res 204(1-2): 60–77. 10.1016/j.heares.2005.01.002.

[29] Breiman L (2001) Random forests. Mach Learn 45(1): 5–32. 10.1023/a:1010933404324.

[30] Dai C, Wood MW, Gan RZ (2007) Tympanometry and laser Doppler interferometry measurements on otitis media with effusion model in human temporal bones. Otol Neurotol 28(4): 551–558. 10.1097/mao.0b013e318033f008.

[31] Morris DP, Wong L, van Wijhe RG, and Bance ML (2006) Effect of adhesion on the acoustic functioning of partial ossicular replacement prostheses in the cadaveric human ear. J Otolaryngol 35(1): 22–25. 10.2310/7070.2005.4116.

[32] Potsangbam DS, Akoijam BA (2019) Endoscopic transcanal autologous cartilage ossiculoplasty. Indian J Otolaryngol Head Neck Surg 71: 54–59. 10.1007/s12070-018-1518-x.

[33] Bahmad F Jr and Merchant SN (2007) Histopathology of ossicular grafts and implants in chronic otitis media. Ann Otol Rhinol Laryngol 116(3): 181–191. 10.1177/000348940711600304.

[34] Quesnel AM, Ishai R, and McKenna MJ (2018) Otosclerosis: Temporal bone pathology. Otolaryngol Clin North Am 51(2): 291–303. 10.1016/j.otc.2017.11.001.

[35] Merchant GR and Neely ST (2023) Conductive hearing loss estimated from wideband acoustic immittance measurements in ears with otitis media with effusion. Ear Hear 44(4): 721–731. 10.1097/AUD.0000000000001317.

[36] Park HW, Ahn J, Kang MW, and Cho Y-S (2020) Postoperative change in wideband absorbance after tympanoplasty in chronic suppurative otitis media. Auris Nasus Larynx 47(2): 215–219. 10.1016/j.anl.2019.08.010.

[37] Asta B, Bozdemir K, Sahin MI (2024) Investigation of ambient-pressure absorbance characteristics of cartilage-grafted tympanic membranes. J Laryngol Otol 138(9): 893–901. 10.1017/S002221512400046X.

